# Temporal transcriptome analysis of the endothelial response to tumour necrosis factor

**DOI:** 10.1101/2023.06.04.543378

**Authors:** EC Struck, T Belova, P Hsieh, J Odeberg, ML Kuijjer, P Dusart, LM Butler

## Abstract

The vascular endothelium acts as a dynamic interface between blood and tissue and provides an anti-inflammatory and anti-thrombotic surface under normal conditions. Tumour necrosis factor-α (TNF), a cytokine that drives acute and chronic inflammation, induces numerous transcriptional changes in endothelial cells (EC). However, the overall temporal dynamics of this response have not been fully elucidated. Here, we conducted an extended time-course analysis of the EC response to TNF, from 30 minutes to 72 hours. We identified TNF-regulated genes and used weighted gene correlation network analysis (WGCNA) to decipher co-expression profiles, uncovering two distinct temporal phases - an acute response initiated between 1-4 hours, followed by a later phase initiated between 12-24 hours. Several previously uncharacterised genes were strongly regulated during the acute phase, while the majority in the later phase were interferon-stimulated genes (ISGs). A lack of interferon transcription indicated that this TNF-induced ISG expression was independent of *de novo* interferon production and autocrine signalling. Furthermore, we observed two different groups of genes whose transcription was inhibited by TNF, those that resolved towards baseline levels over time, and those that did not. Our study provides insights into the global temporal dynamics of the EC transcriptional response to TNF, highlighting distinct gene expression patterns during the acute and later phases. These findings may be useful in understanding the role of EC in inflammation and developing TNF signalling-targeting therapies.

## INTRODUCTION

The vascular endothelium is a dynamic interface between blood and tissue that has a role in the regulation of coagulation, blood pressure, solute movement, and inflammation. The resting endothelium is an anti-inflammatory and anti-thrombotic surface, which is unreceptive to interactions with circulating blood cells (Ley and Reutershan, 2006; Yau et al., 2015). The cytokine tumour necrosis factor-α (TNF) is a key driver of acute and chronic inflammation (Jang et al., 2021; Webster and Vucic, 2020) and can bind to endothelial cells (EC) via TNF-receptors 1 and 2. This interaction induces signalling cascades that regulate the activity of several transcription factors, including NF-kappaB (NFκB) and activator protein-1 and 2 (Baud and Karin, 2001; Vandenabeele et al., 1995), leading to various cellular responses, such as the expression of adhesion molecules and chemokines that facilitate leukocyte recruitment into tissue (Liao, 2013). TNF can also induce the EC expression of interferon regulatory factors and interferon-stimulated genes (ISGs) (Venkatesh et al., 2013; Yan et al., 2017). Indeed, the concept that EC are multifaceted conditional innate immune cells has gained traction in recent years (Lu et al., 2022; Shao et al., 2020). However, the global dynamics of the EC response to TNF is not well understood, with existing studies tending to focus on specific gene(s) or phenotypic characteristics, e.g., (Brandt et al., 2022; Jung et al., 2020; Ulfhammer et al., 2006), or global transcriptional changes at a single, or small number, of time points (Rastogi et al., 2012; Ryan et al., 2022). The same is true of studies of the TNF response in other cell types, such as the immortalised embryonic kidney cell line HEK293 (Bouwmeester et al., 2004; Ma et al., 2009) or fibroblasts (Hao and Baltimore, 2013; Paulsen et al., 2013). Existing studies also tend to neglect the influence of chromosomal sex on cell behaviour (Lu et al., 2018), despite reported differences between male and female EC, e.g., in preeclampsia (Zhou et al., 2019), and following exposure to shear stress (James and Allen, 2021), X-ray induced damage (Campesi et al., 2022) and hyperoxia (Zhang and Lingappan, 2017).

Here, we profiled the EC transcriptional response to TNF over an extended time, including 11 time points ranging from 30 minutes to 72 hours post stimulation. We identified TNF up regulated or down regulated genes and used weighted gene network correlation analysis to decipher co-expression profiles, revealing two distinct temporal phases; an acute response initiated between 1-4 hours, and a subsequent later one between 12-24 hours. Sex-based analysis revealed a high similarity in response profile between female and male EC. Several completely uncharacterised genes were strongly regulated during the acute phase of the response, while the majority of those in the latter phase were interferon stimulated genes (ISG). Our data indicated that TNF-induced ISG expression was independent of *de novo* interferon production and subsequent autocrine signalling. All data is available on (https://butlerlab.shinyapps.io/temporal_TNF_response/), which users can view on a gene centric or regulation-profile basis.

## RESULTS

Existing studies of EC responses to inflammatory stimuli, such as TNF, focus primarily on early signalling and associated transcriptional changes. Here, we profiled global changes in the EC transcriptome following TNF stimulation, over an extended time course. Human umbilical vein endothelial cells were extracted and pooled by donor sex, before treatment with or without TNF for 0.5, 1, 2, 4, 6, 8, 12, 24, 36, 48 or 72 hours, and subsequent analysis by RNA sequencing (Figure S 1A).

### TNF induces EC transcriptome modifications over an extended time course

Average DESeq2 expression values for each treated sample (biological replicates: n=3 male and n=2 female) were normalised to the sex- and time-matched untreated controls, to identify differentially expressed genes (DEG). 2099 positive and 2162 negative DEG were identified (fold change vs. untreated control FC log2 > 1 and FC log2 < -1, respectively [in both male and female sample sets], TPM>1 at least one sample time point, raw counts >10 in all samples, adjusted by p-value) (Figure S1 B and methods for analysis details) (Table S1, Tabs 1-2). Of these, a total of 918 genes were further classified as up regulated in response to TNF (Figure 1 A.i), and 210 as down regulated (Figure 1B.i), based on a TNF-induced change from a relatively stable baseline expression in untreated (control) EC over time (Figure S1 C.i and D.i). Genes classified as DEG due to changes in expression in untreated control EC over time (Figure S1 C.ii and S1 D.ii), or a TNF-induced lag in such changes (Figure S1 C.iii), were excluded, as such changes may be linked to *in vitro* culture conditions and/or the influence of TNF on other inflammation-independent temporal processes.

**Figure 1.**
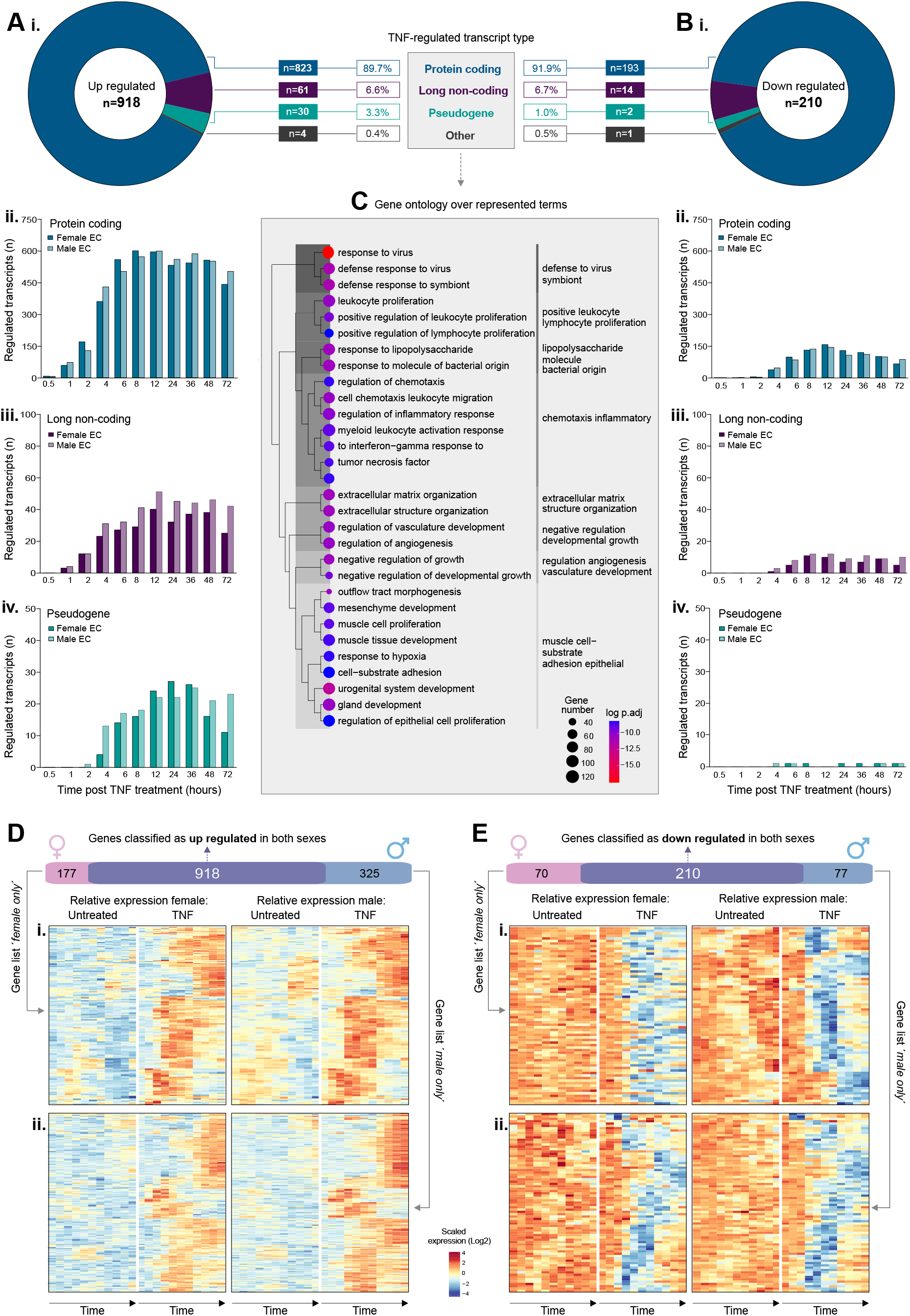
Overview of tumour necrosis factor (TNF) regulated genes in human endothelial cells. Human umbilical vein endothelial cells (EC, n=5) were treated with or without tumour necrosis factor alpha (TNF) and harvested at 0.5, 1, 2, 4, 6, 8, 12, 24, 36, 48 or 72 hrs, before RNAseq analysis. Genes were classified as (**A**) up regulated or (**B**) down regulated by TNF (fold change vs. untreated control >2 or <0.5, respectively). Plots show: (i) total TNF regulated genes (in both sexes) and corresponding biotype, and number of (ii) protein coding, (iii) lncRNA or (iv) pseudogenes regulated at each time point, by sex. (**C**) Gene ontology analysis showing over-represented terms for all TNF regulated transcripts. (**D**) and (**E**) show the number of genes classified as either up regulated or down regulated, respectively, in females or males only, or in both sexes. Heatmaps show relative expression in control or TNF treated female (left) or male (right) samples for genes reaching the threshold for classification as TNF-regulated in (i) males or (ii) females only.

The majority of up regulated genes were classified as protein coding (845/952; 88.7 %), followed by long non-coding RNAs (lncRNA) (69/925; 7.2%), pseudogenes (34/952; 3.8%) and ‘other’ (3 TEC, 1 sRNA, 1 miscellaneous RNA) (Figure 1A.i). A similar profile was observed for down regulated transcripts, with the majority classified as protein coding (210/232; 90.5 %), followed by lncRNA (16/232; 6.9%), pseudogenes (4/232; 1.7%) and ‘other’ (1 TEC, 1 sRNA) (Figure 1B.i). We performed gene ontology analysis (Ashburner et al., 2000b) to identify over-represented groups in genes classified as regulated by TNF; significant enrichment terms included ‘*response to cytokine*’ (FDR 6.0 ×10^-30^) and ‘*immune system processes*’ (FDR 5.3 ×10^-29^) (Figure 1C) (Table S2, Tab 1).

To investigate the temporal profile of genes up or down regulated in response to TNF, we determined the number of transcripts classified as such at each time point, in each sex (Figure 1 A.ii-iv and B.ii-iv). A limited number of protein coding transcripts (n=10) were classified as up regulated already by 30 minutes post-stimulation, in both sexes (Figure 1 A.ii), including those encoding for components of the NFκB-signalling pathway (*NFKBIA*, *NFKBIZ*), chemokines regulated by this pathway (*CXCL2*, *CXCL3*, *IL6*) and *FAM167A*, which was recently described as an activator of the non-canonical NFκB-pathway in chronic myeloid leukaemia (Yang et al., 2022) (Table S2, Tab 1).

A small panel of lncRNAs were classified as up regulated by 1-hour post-stimulation, in both sexes (n=10) (Figure 1 A.iii); including those known to be expressed in response to inflammation in EC, e.g., *MIR155HG* (Barros Ferreira et al., 2022), but also those not previously described in this context, e.g., wound and keratinocyte migration-associated long noncoding RNA 2 (*WAKMAR2*), which restricts inflammatory chemokine production in keratinocytes, and enhances their migratory capacity (Herter et al., 2019).

TNF-responsive pseudogenes tended to be classified as up regulated slightly later in the time course than protein coding or long non-coding genes. From a total of 30 pseudogenes classified as up regulated by TNF (Figure 1 A.iv), 10/30 [33%] were members of the ferritin gene family (*FTH1P2, 7, 8, 10, 11, 12, 15, 16, 20, 23*), some of which have known regulatory functions (Di Sanzo et al., 2020). Ferritin has a role in EC angiogenesis (Tesfay et al., 2012) and chemokine signalling (Li et al., 2006); a potential role of this family in the EC response to inflammatory stimulation remains to be explored. Overall, the total number of genes that were up regulated in response to TNF was highest between 12- and 24-hours post-stimulation (Figure 1 A.ii, iii and iv).

In contrast to TNF-induced up regulated genes, no protein coding genes were classified as down regulated at the earliest time point (Figure 1 B.ii). Those classified as such within the first four hours (n=54), included *HOXA9*, which inhibits NFκB dependent EC activation (Trivedi et al., 2007). The total number of down regulated protein coding transcripts peaked around 12 hours (Figure 1 B.ii). lncRNAs classified as down regulated in response to TNF were also classified as such later than those that were up regulated (Figure 1B.iv), and only a small number of pseudogenes were consistently down regulated across the time course (Figure 1 B.iv).

Thus, the global primary EC response to TNF stimulation predominantly consists of the induction of protein coding and long non-coding gene expression.

### TNF-induced changes are comparable between male and female endothelial cells

Differences in inflammatory response have been previously described in female and male EC (Addis et al., 2014; Zhou *et al*., 2019). In our analysis described above, 177 genes were classified as up regulated only in female samples and 325 only in male samples (Figure 1 D), and 70 genes were classified as down regulated only in female samples and 77 only in male samples (Figure 1 E) (Table S1, Tab 3). Heatmap plots of genes classified as up or down regulated only in female EC (Figure 1 D.i and E.i, respectively) or only in male EC (Figure 1 D.ii and E.ii, respectively), revealed similar patterns of regulation over the time course in the other respective sex (relative expression in female EC on the left and male EC on the right). Thus, these differences in classification were likely due to the strict thresholding criteria we initially applied to identify the most consistently and strongly regulated genes, rather than a fundamental difference between the responsiveness of EC from each sex. Indeed, when we applied a threshold of up-or down regulation in one or more samples of one sex (FC log2 abs >1), versus no regulation in any samples of the other sex (FC abs <1.1 [FC log2 <0.1375]) no genes were classified as sex-specifically regulated by TNF. Multidimensional scaling revealed high global similarity in TNF-induced transcriptome modifications over the time course in female (Figure S2 A.i) and male (Figure S2 A.ii) EC. Thus, the chromosomal composition of the EC does not appear to markedly effect the global transcriptional response to TNF stimulation.

To identify baseline (unstimulated) differences in gene expression in female versus male EC, we performed differential expression analysis between the control unstimulated samples of each sex. 99 genes were classified as differentially expressed between the sexes at every time point (FC log2 abs >1, adjusted p<0.05), with 58 genes being higher in females and 41 higher in males (Figure S2 B). As expected, Y chromosome genes represented the most significantly differentially expressed genes in male EC (Figure S2 B.i) and the long noncoding RNA *XIST*, a regulator for X-inactivation (Loda and Heard, 2019), was the most significantly differentially expressed gene in female EC (Table S2, Tab 3).

### Weighted gene network analysis reveals TNF-induced gene signatures

To explore the potential relationship between TNF-regulated genes, in terms of expression dynamics over time, we performed a weighted co-expression gene network analysis (WGNCA) (Langfelder and Horvath, 2008), where correlation coefficients between all transcripts across the sample set (male and female samples were handled together) were calculated and subsequently clustered into 48 related modules (Figure 2 A.i), based on expression profile similarity. In addition to the identification of co-regulated genes, this analysis could potentially highlight genes with a currently unknown roles in specific stages of the EC response to TNF.

**Figure 2.**
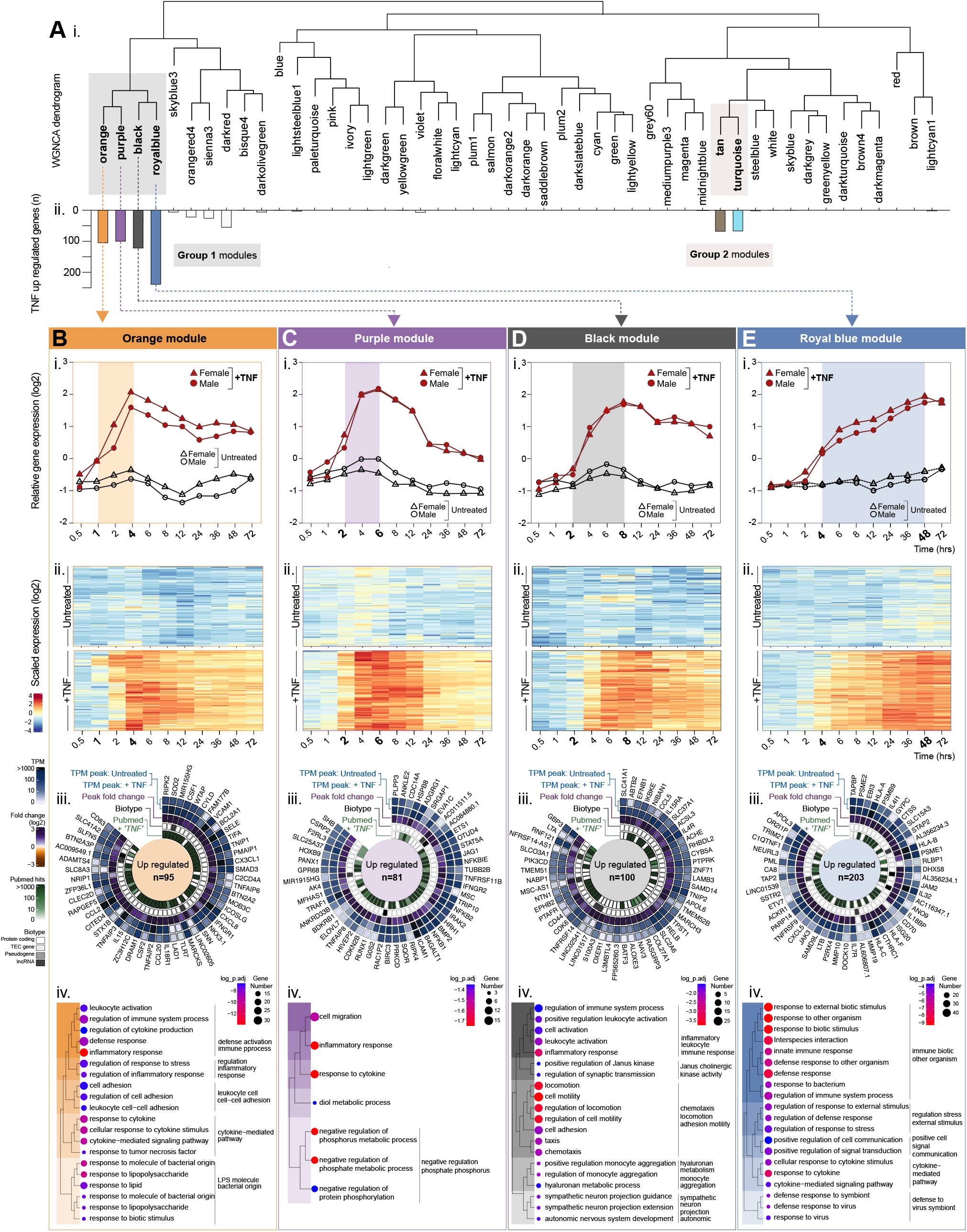
Weighted network correlation analysis (WGNCA) reveals temporal relationships between TNF up regulated genes: Group 1 ‘early induced’. Human umbilical vein endothelial cells (EC, n=5) were treated with or without tumour necrosis factor alpha (TNF) and harvested at 0.5, 1, 2, 4, 6, 8, 12, 24, 36, 48 or 72 hrs, before RNAseq analysis. Weighted correlation network analysis (WGCNA) was used to cluster genes into modules, based on expression pattern similarity across sample sets. (**A**) (i) Dendrogram showing WGCNA modules and (ii) corresponding distribution of genes previously classified as TNF up regulated. For modules (**B**) *orange*, (**C**) *purple*, (**D**) *black*, (**E**) *royal blue*: (i) relative gene expression plots displaying the module eigengenes and (ii) heatmaps showing the temporal expression profile for all genes in the module. (iii) Circle plots for the top 50 genes classified as TNF-upregulated within each module: showing expression values in control and stimulated EC, the peak fold change, the biotype and number of PubMed hits for ‘*gene name’* + ‘TNF’, and (iv) over-represented terms by gene-ontology analysis (biological processes).

#### TNF up regulated genes have two main regulation profiles

The majority of genes we earlier classified as up regulated in response to TNF stimulation fell into two main module regions on the WGNCA dendrogram (Figure 2 A.i), occupying neighbouring leaves on common clades: annotated as group 1 [*orange*, *purple*, *black* and *royal blue*] and group 2 [*tan* and *turquoise*] modules (Figure 2 A.ii) (see shaded boxes).

### Up regulated group 1-Early induction (1-4 hours post stimulation)

Group 1 modules contained genes that were up regulated by 1 hour [*orange*] (Figure 2 B.i-ii), 2 hours [*purple*, *black*] (Figure 2 Ci-ii and Di-ii) or 4 hours [*royal blue*] (Figure 2 E.i-ii) post-TNF stimulation. Genes in the *orange* (Figure 2 B.i-ii), *purple* (Figure 2 C.i-ii) and *black* (Figure 2 D.-ii) modules reached a peak between 4 and 8 hours, after which the differential expression vs. control gradually declined. Several genes with well-established roles in the initial stages of inflammation were in the ‘earliest responder’ *orange* module (see Figure 2B.iii for the top 50 up regulated genes with highest correlation to the eigengene), including those encoding for components of the NFκB-signalling pathway e.g., *NFKBIA*, *NFKBIZ*, leukocyte adhesion receptors e.g., *VCAM1, SELE*, and various chemokines or cytokines, e.g., *CXCL8*, *CX3CL1*, *CCL2* and *CCL20*. A PubMed search, retrieving the number of publications containing the search terms [gene] and [TNF], revealed most of the *orange* module top 50 genes had been previously referred to in the context of the TNF response (Figure 2 B.iii). However, some, e.g., *FAM177B*, an - uncharacterised gene with little to no expression under baseline conditions, have no previous reported link with the EC response to TNF, or indeed inflammation processes in general. This was also true for some genes in the other modules, e.g., *purple* module long non-coding gene *MIR1915HG*, which currently has no assigned function, and *black* module gene *ALOXE3*, which encodes for a member of the lipoxygenase family that was recently reported as expressed in EC and up regulated by shear stress exposure (Sabbir et al., 2022). One could speculate that such genes encode for proteins with currently unknown important roles in the initial stages of inflammation, and thus represent interesting candidates for functional investigation. Gene ontology analysis of TNF up regulated genes in the *orange* (Figure 2 B.iv), *purple* (Figure 2 C.iv) and *black* (Figure 2 D.iv), revealed over representation of similar inflammation-related terms, such as *inflammatory response* and *response to cytokine* (Table S2, Tab 4, A-C).

In contrast to the other group 1 modules, TNF up regulated genes in the *royal blue* module remained elevated across the 72-hour time course (Figure 2 E.i-ii) and contained genes encoding for several types of pattern recognition receptors (PRR), e.g., toll-like- (*TLR2, TLR5)*, RIG-I-like- *(DDX58*, *DHX58*, *IFIH1)* and NOD-like- (*NOD2*, *NLRC5*) receptors, and cyclic guanosine monophosphate adenosine monophosphate synthase (*cGAS*). This module also contained *ISG20,* an interferon stimulated gene (ISG) that encodes a nuclease enzyme that can cleave viral RNA. Gene ontology analysis of TNF up regulated genes in this module revealed that overrepresented terms included those linked to viral defence, such as *response to virus* (Figure 2 E.iv) (Table S2, Tab 4, D).

### Up regulated group 2 - Delayed induction (12-24 hours post stimulation)

Group 2 modules [*turquoise* and *tan*] (Figure 3 A) contained genes that were up regulated between 12-24 hours post-TNF treatment, with the highest differential expression vs. control at 72-hours (Figure 3 Bi-ii and Ci-ii). TNF-up regulated genes in the *turquoise* module (Figure 3 B) included those encoding for a panel of interferon-induced cytokine ligands for the antigen presenting cell receptor CXCR3: *CXCL9*, *CXCL10* and *CXCL11* (Figure 3 B.iii), the expression of which is linked with viral infection, progression and replication control (Callahan et al., 2021; Karin, 2020; Yin et al., 2019). Other ISGs in this module included *IFIT2*, *IFIT5* and *IFIT35*, that, to our knowledge, have not previously been reported as TNF-regulated in EC. Gene ontology analysis of up regulated genes in the *turquoise* module revealed over representation of terms associated with *defence to virus symbiont* (Figure 2 B.iv), in addition to general inflammation terms, such as *response to cytokine* (Table S2, Tab 4, E). TNF-up regulated genes in the *tan* module also included a large panel of ISGs e.g., among others *IFIT1*, *IFI6*, *IFI27*, *OAS1/2/3/L*, *MX1/2* and *IFITM1* (Figure 3 C.iii) and, correspondingly, gene ontology analysis revealed a highly significant over representation of terms related to response to interferon and viral infection (Figure 3 C.iv) (Table S2, Tab 4, F). Thus, this latter stage of the TNF response was dominated by the induction of interferon/anti-viral related gene transcription. Indeed, a PubMed search for studies citing TNF-up regulated genes in the *turquoise* (Figure 3D) and *tan* (Figure 3E) modules, together with the term ‘interferon’, revealed most had been previously reported in this context (40/59 [68%] and 57/66 [88%], respectively). However, only a small proportion also included the term ‘endotheliaĺ (Figure 3 D and E, dark shaded bars), indicating that these pathways are less well understood in this cellular context. Genes in the *turquoise* module with only one, or no hits linking them to interferon signalling (Figure 3D), included several non-coding genes, which are typically less well studied than protein coding genes, e.g., *HCP5,* which has polymorphisms linked to HIV viral load (Thorner et al., 2016) and is a susceptibility locus for Kawasaki disease, a systemic vasculitis of infants and children (Kim et al., 2017), the uncharacterised pseudogene *AKRD26P1,* and *LINC01094* (Figure 3 D, right panels). Thus, it is possible that such genes have currently unknown roles in EC interferon-related signalling, based on the similarity of their temporal expression profile with others in the module.

**Figure 3.**
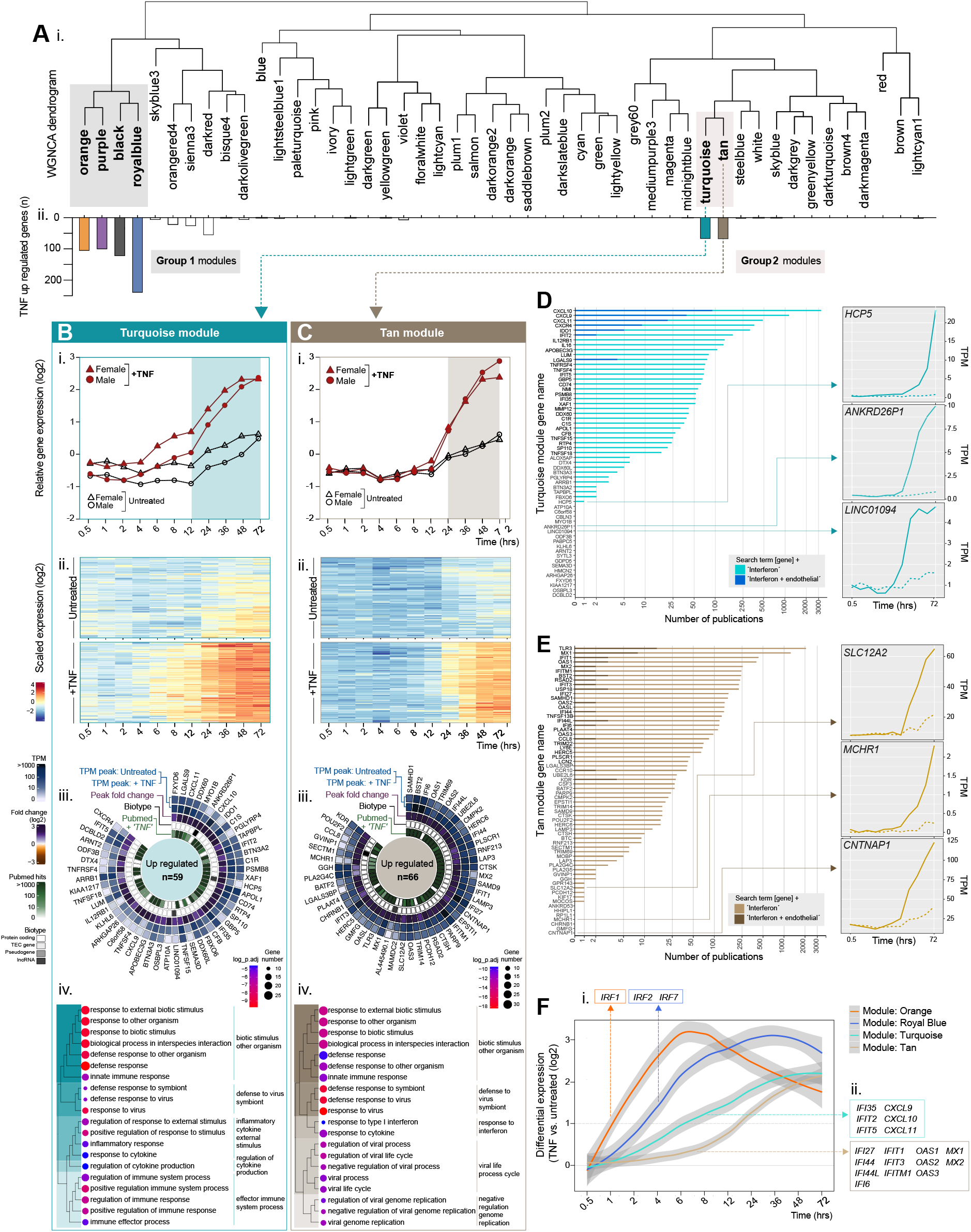
Weighted network correlation analysis (WGNCA) reveals temporal relationships between TNF up regulated genes: Group 2 ‘delayed induced’. Human umbilical vein endothelial cells (EC, n=5) were treated with or without tumour necrosis factor alpha (TNF) and harvested at 0.5, 1, 2, 4, 6, 8, 12, 24, 36, 48 or 72 hrs, before RNAseq analysis. Weighted correlation network analysis (WGCNA) was used to cluster genes into modules, based on expression pattern similarity across sample sets. (**A**) (i) Dendrogram showing WGCNA modules and (ii) corresponding distribution of genes previously classified as TNF down regulated. For modules (**B**) *tan* and (**C**) *turquoise*: (i) relative gene expression plots displaying the module eigengenes and (ii) heatmaps showing the temporal expression profile for all genes in the module. (iii) Circle plots for the top 50 genes classified as TNF-up regulated within each module: showing expression values in control and stimulated EC, the peak fold change, the biotype and number of PubMed hits for ‘*gene name’* + ‘TNF’, and (iv) over-represented terms by gene-ontology analysis (biological processes). (**D**) and (**E**) show the number of hits returned for TNF up regulated genes in the *tan* or *turquoise* modules, respectively, in a PubMed search for ’*gene name*’ + ’interferon’ and ’*gene name*’ + ’interferon’ + ’endothelial’, with temporal expression plots for selected examples (created using the website tool provided as part of this study). (**F**) Temporal distribution of interferon-related genes upregulated by TNF across *orange*, *royal blue*, *tan*, and *turquoise* modules.

11 genes in the *tan* module had only one, or no hits linking them to interferon signalling, including the solute transported *SLC12A2*, the G-protein coupled receptor *MCHR1* and the paranodal junction component *CNTNAP1* (Figure 3 E, right panel). The role of these genes in inflammation, and any possible connection to EC interferon and/or viral response signalling remains to be established.

### TNF-induced expression of ISGs is not driven by *de novo* interferon production

Although the mechanisms of TNF-induced ISG expression in EC are not well understood, previous studies have reported that it is driven by *de novo* production and subsequent autocrine signalling of type I interferon (IFNβ) (Venkatesh *et al*., 2013). We found that three of the nine members of the interferon regulatory factor family (IRF1-9), which are critical for the induction of type I interferon (McNab et al., 2015), were up regulated by TNF, and all were found in the group 1 ‘early respondinǵ modules *orange* (*IRF1*) and *royal blue* (*IRF2* and *IRF7*) (Figure 3 F.i), thus temporally preceding the induction of the majority of the ISGs, which fall in modules *turquoise* and *tan* (Figure 3 F.ii). However, only a modest TNF-induction of *IFNB1* was observed at later time points (max. any sample, any time point = 1.02 TPM [24h]) (Table S4, Tab 5); importantly, TNF-induced expression of many ISGs preceded the time point at which *IFNB1* was expressed at detectable levels e.g., *ISG20* (Figure 4 A.i), *IFIT3* (Figure 4 A.ii), *IFI35* (Figure 4 A.iii), *CXCL10* (Figure 4 A.iv) *CXCL11* (Figure 4 A.v), and *MX1* (Figure 4 A.vi). The same was also observed for various pattern recognition receptors, e.g., *DDX58* (Figure 4 A.vii), *TLR2* (Figure 4 A.viii), *NOD2* (Figure 4 A.ix), and *IFIH1* (Figure 4 A.x). Thus, in this system, the transcriptional induction of such genes was not driven by autocrine IFNβ signalling. TNF-induced changes in the expression of genes encoding for key components of the interferon/anti-viral response pathways (JAK/STAT, NFκB-IRF1, TLR, RLR and cGAS/STING) are summarised in Figure 4 B.

**Figure 4.**
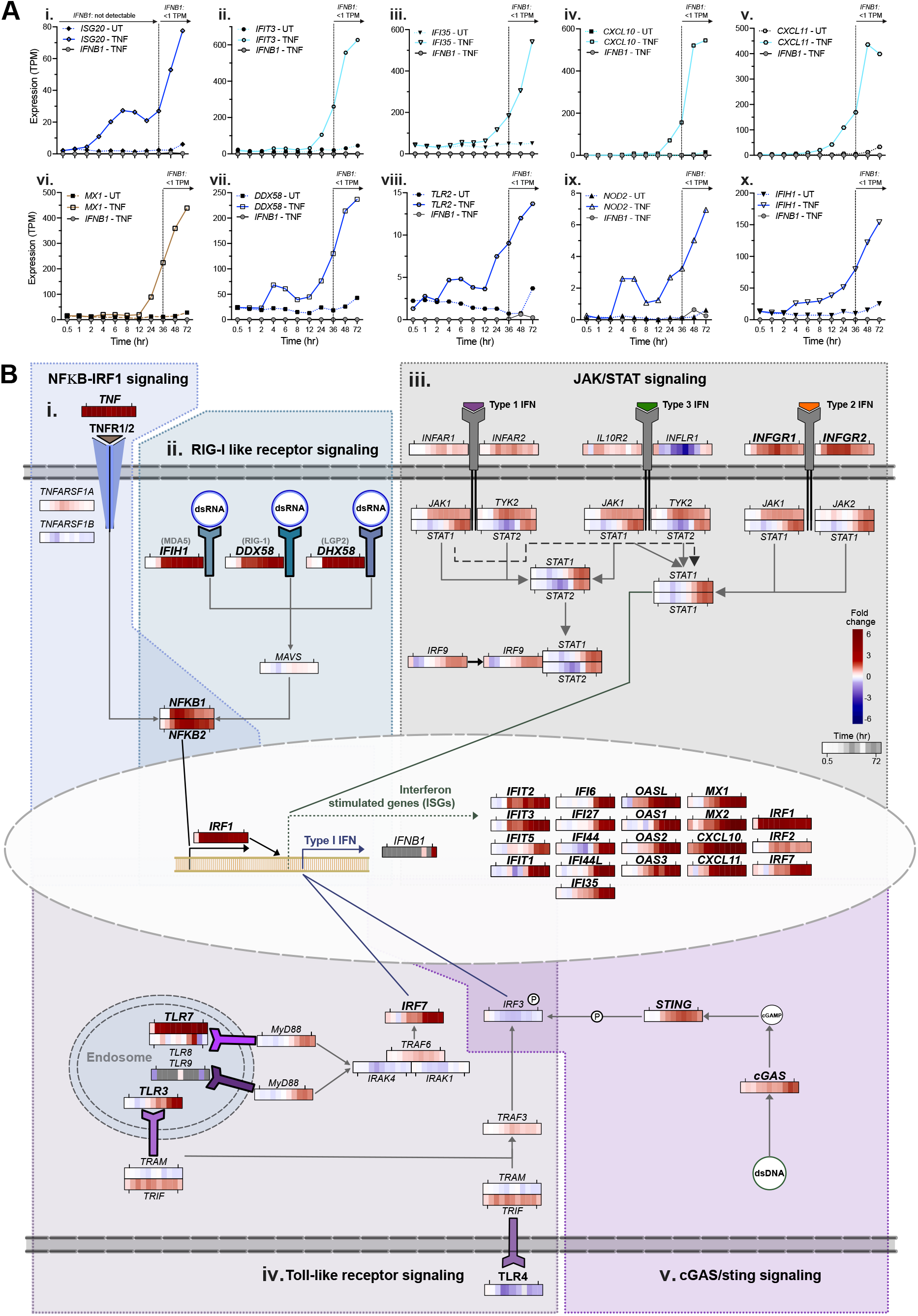
TNF up regulation of interferon-stimulated and pattern recognition receptor gene expression is independent of *de novo* interferon production. (**A**) Temporal expression profiles in unstimulated control or tumour necrosis factor alpha (TNF) stimulated EC for *IFNB1* and (i) *ISG20*, (ii) *IFIT3*, (iii) *IFI35*, (iv) *CXCL10*, (v) *CXCL11*, (vi) *MX1*, (vii) *DDX58*, (viii) *TRL2*, (ix) *NOD2* and (x) *IFIH1* (sample set F). (B) Summary of key genes and pathways linked to interferon stimulated gene expression: (i) NFKB-IRF1 signalling (adapted from Feng et al., 2021), (ii) RIG-I like receptor signalling (adapted from Rehwinkel and Gack., 2020), (iii) JAK-STAT signalling (adapted from Schneider et al., 2014), (iv) TLR-signalling (adapted from Duan et al., 2022), and (v) CGAS-Sting signalling (adapted from Feng et al., 2021). Heatmaps show the differential gene expression between unstimulated control and TNF stimulated EC for the adjacent gene. Grey squares in the heatmap are the result of zero TPM values, thus differential expression is not calculated. Bold gene symbols denote those that were classified as TNF upregulated. Heatmaps were created using the website tool provided as part of this study.

#### TNF down regulated genes have two main regulation profiles

The majority of genes classified as down regulated in response to TNF stimulation fell into two main regions on the WGNCA dendrogram (Figure 5 A.i), occupying neighbouring leaves on common clades: annotated as group 1 [*dark orange*, *saddle brown*] and group 2 [*green* and *light yellow*] modules (Figure 5 A.ii) (highlighted with shaded boxes). Modules in both groups contained genes that were down regulated by TNF between 2- and 4-hours post-stimulation (Figure 5 Bi-ii, Ci-ii, Di-ii, and Ei-ii).

**Figure 5.**
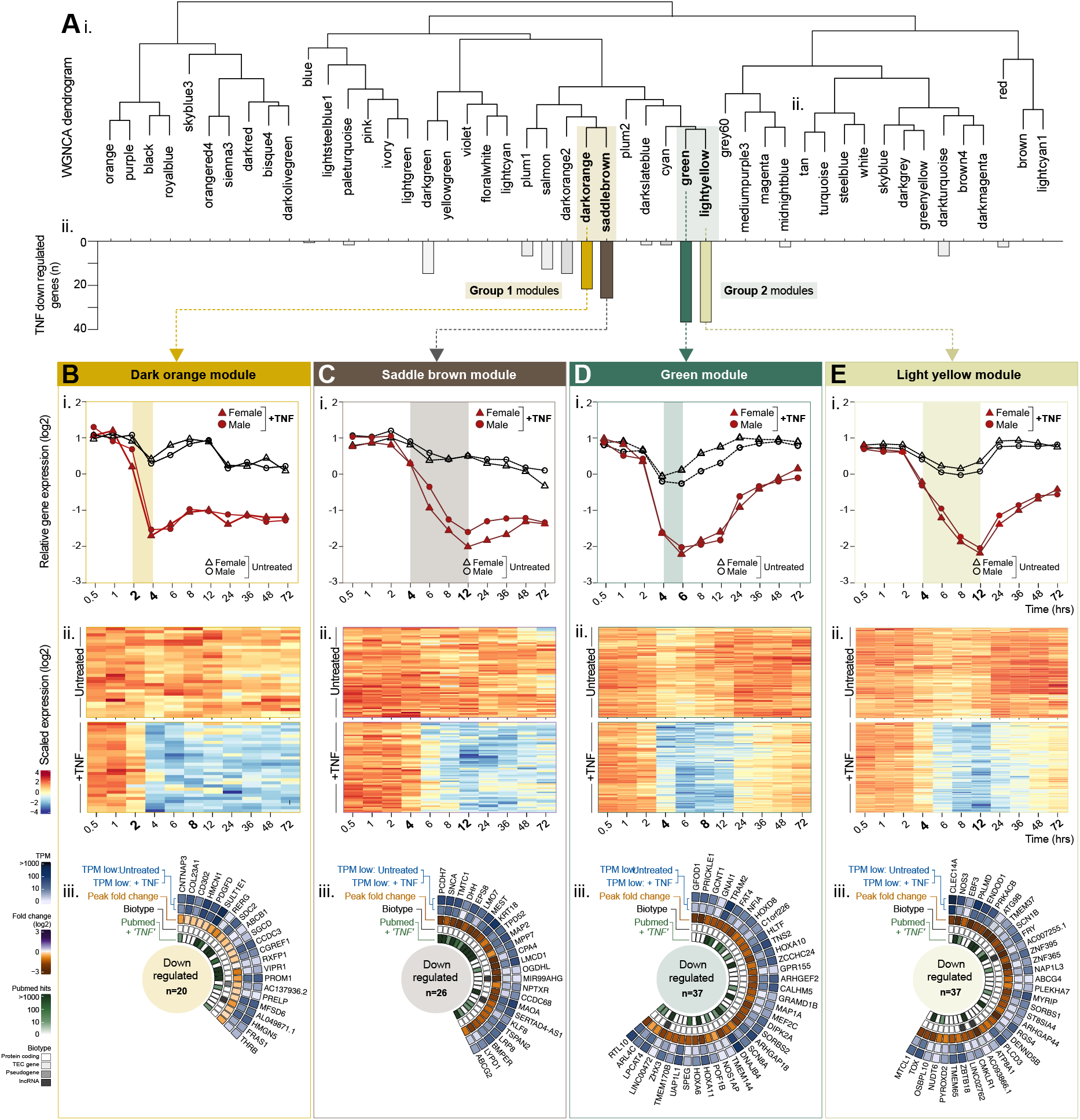
Weighted network correlation analysis (WGNCA) reveals temporal relationships between TNF down regulated genes. Human umbilical vein endothelial cells (EC, n=5) were treated with or without tumour necrosis factor alpha (TNF) and harvested at 0.5, 1, 2, 4, 6, 8, 12, 24, 36, 48 or 72 hrs, before RNAseq analysis. Weighted correlation network analysis (WGCNA) was used to cluster genes into modules, based on expression pattern similarity across sample sets. (**A**) (i) Dendrogram showing WGCNA modules and (ii) corresponding distribution of genes previously classified as TNF down regulated. For modules (**B**) dark orange, (**C**) saddle brown, (**D**) green, (**E)** light yellow: (i) relative gene expression plots displaying the module eigengenes and (ii) heatmaps showing the temporal expression profile for all genes in the module. (iii) Circle plots for genes classified as TNF-down regulated within each module: showing expression values in control and stimulated EC, the peak fold change, the biotype and number of PubMed hits for ‘*gene name’* + ‘TNF’.

### Down regulated group 1 – Inhibition maintained over time

Genes in the group 1 modules *dark orange* (Figure 5 B.i-ii) and *saddle brown* (Figure 5 C.i-ii) reached maximum down regulation 4- and 12-hours post stimulation, respectively, an effect that was maintained across the remainder of the time course. There were no significantly enriched gene ontology terms in lists of down regulated genes appearing in the group 1 down regulated modules, possibly due to the low numbers, or lack of previous reports of gene function.

### Down regulated group 2 - Inhibition resolved over time

In contrast to group 1, TNF down regulated genes in group 2 modules, *green* (Figure 5 D.i-ii) and *light yellow* (Figure 5 E.i-ii), trended back towards baseline level after reaching maximum down regulation at 6- and 12-hours post stimulation, respectively. Again, there were no significantly enriched gene ontology terms in any of lists of down regulated genes appearing in the highlighted modules; indeed, an automated PubMed search for the search terms [down regulated gene] and [TNF], retrieved markedly fewer hits than the equivalent search for genes we previously classified as up regulated by TNF (Figure 5 B-E.iii). Despite a lack of significant enrichment terms, some links between constituent genes could be observed e.g., four out of five TNF down regulated homeobox family transcription factor genes were classified into module *green* (*HOXA6*, *HOXA10, HOXA11* and *HOXD8*) (Figure 5 D.iii). Whilst *HO×10* has been reported as an activator of canonical NF-κB signalling in pancreatic cancer cells (Li et al., 2022), and the antisense to *HOXA11* (*HOXA11-AS*) linked to protection of EC barrier function following injury (Yuan et al., 2022), insight into the potential role of these genes in the EC TNF response is currently lacking.

In both TNF down regulated modules group 1 and 2, gene baseline expression levels and TNF-response profiles were similar between male and female samples (Figure 5 B.i-E.i and Figure S2 C).

### Visualisation of temporal gene regulation pathways using the website tool

We have created a website resource that allows users to perform both gene centric or module-based lookup of our endothelial TNF time course data. Key features include a data viewer, to observe the transcriptional responses of specific genes (Figure S3 A), or the WGCNA module into which they were classified (Figure S3 C), and the generation of vector-image downloadable expression plots for both predefined and custom gene lists, e.g., TNF regulated CXCL- and CCL-chemokines (Figure S3 B). The dataset can also be analysed to identify genes with high correlation across conditions with any given input gene (Figure S3 D).

## DISCUSSION

Here, we used RNA sequencing to measure the temporal response of EC to TNF stimulation, incorporating 11 time points up to 72h. Following the identification of TNF regulated genes, we used weighted network correlation analysis to understand the global temporal context of these transcriptomic changes. To our knowledge, this is the first study to map the EC TNF response at a transcriptome-wide level in such temporal detail.

We identified two main profiles into which TNF-induced genes could be classified - those with an ‘earlý or ‘delayed’ induction. Early induced genes (∼1-4 hours post stimulation) included many previously well studied in this context, such as those encoding for EC leukocyte adhesion receptors (e.g., *SELE*, *VCAM1* and *ICAM1*) (Pober, 2002), but we also identified several genes with similar expression dynamics that encoded for completely uncharacterised proteins (e.g.. *FAM177B*), which could be interesting candidates for future study in the context of inflammation. TNF up regulated genes with a delayed induction (12-24 hours post stimulation) were primarily ISGs, whose regulation by inflammatory cytokines in EC is generally not well understood, but our observations were consistent with one recent study which reported the induction of a late stage interferon response in EC, following TNF stimulation (Valenzuela, 2022). ISG expression is primarily considered to be driven by the production and subsequent signalling of interferon, via canonical (JAK-STAT) or non-canonical pathways (Mazewski et al., 2020). We found that three out of the nine members of the interferon regulatory factor family (*IRF1*, *2* and *7*), which are critical for the induction of interferon (McNab *et al*., 2015), were up regulated by TNF in EC at time points that preceded ISG expression. Of these, *IRF1* and *IRF7* have been implicated as positive regulators of type I interferon production (Honda et al., 2006) and previous reports have shown that TNF-induced expression of ISGs, such as *CXCL9* and *CXCL10*, in murine EC was dependent on *IRF1*-induced *de novo* production of IFNβ (*IFNB1*), and its subsequent autocrine signalling through STAT1 (Venkatesh *et al*., 2013). However, our data indicated that EC TNF-induced expression of ISGs was independent of *de novo* production of interferon, as we did not observe its transcription prior to ISG expression.

We observed an up regulation of genes encoding for cyclic GMP–AMP (CGAS) and the cyclic GMP–AMP receptor stimulator of interferon genes (STING), a system which detects pathogenic DNA (Hopfner and Hornung, 2020). A recent study showed that the expression of various ISGs that were induced by TNF in fibroblasts, including *CXCL10*, *IFIT1* and *IFIT44* (all of which were also up regulated by TNF in the current study) was markedly reduced in CGAS and STING knockout cells (Willemsen et al., 2021). TNF-dependent mitochondrial damage and mtDNA leakage was shown to underlie this response; one could speculate a similar mechanism contributes occurs in EC.

We observed an up regulation of genes encoding for other pattern recognition receptors, including toll-like- (*TLR2, TLR5)*, RIG-I-like- *(DDX58*, *DHX58*, *IFIH1)* and NOD-like- (*NOD2*, *NLRC5*) receptors. Whilst these receptors are known to induce production of interferon and subsequent expression of ISG in response to bacterial or viral ligands, (Opitz et al., 2009; Uematsu and Akira, 2007), whether or not they have a role in the induction of ISG in EC following TNF production, similar to that reported for CGAS and STING (Willemsen *et al*., 2021), remains to be explored.

We identified several non-coding RNAs within the gene modules that otherwise predominantly contained ISGs, including ENSG00000225886 (antisense to *IFI6*), *NRIR*, a negative regulator of SARS-CoV-2 infection (Enguita et al., 2022) and *LINC02056*, an interferon-inducible transcript with a proposed role in IRF3 nuclear translocation (Xu et al., 2021). As is often the case with non-coding genes, functional annotation of others was lacking e.g., *LINC02051* and *LINC02068*; these are potentially interesting candidates to study in the context of the EC interferon response. Overall, deciphering the relative contributions of various pathways in the TNF-induced expression of ISGs is complex, with potential differences between cell types and species.

We identified a panel of 210 gene transcripts that were at lower levels following TNF-stimulation. Whilst studies of the TNF response tend to focus on genes whose expression in increased in response to stimulation, several of those we identified as down regulated had been previously reported as such, e.g., *NOS3*, the mRNA stability of which is inhibited by TNF (Yan et al., 2008), *DHH*, which prevents EC activation (Chapouly et al., 2020), and *RGS4*, which regulates the secretion of VWF (Patella and Cutler, 2020). However, many had not been previously reported in this context, e.g., *CNR1*, *SLC7A8* and *CLEC14A*, which were amongst the most down regulated by TNF.

Sex differences have been reported in several inflammatory conditions of the vasculature, such as cardiovascular disease (Gao et al., 2019) and thrombosis (Nordstrom and Weiss, 2008). Whilst our data indicates that chromosomal composition alone does not markedly affect the EC response to TNF, a multitude of other factors influence vascular responses *in vivo*, such as sex hormones, which may drive sex linked inflammatory differences (Pabbidi et al., 2018; Rathod et al., 2017).

### Study strengths and limitations

One of the main strengths of our study is size of the dataset generated; we analysed 130 samples, incorporating 35 biological replicates. The global EC transcriptome was analysed at 11 different time points post-TNF treatment, and the inclusion of matched control samples for each sample set, at every time point, allowed us to control for baseline transcriptional changes, such as those due to changes in the microenvironment (Majewska et al., 2021) or cell density (Hamada et al., 2014), which could otherwise be incorrectly annotated as TNF driven. To our knowledge, the website resource we provide (https://butlerlab.shinyapps.io/temporal_TNF_response/) is the most extensive of its type; all data is accessible without the need for bioinformatic expertise. The EC we used in our study were isolated from human umbilical veins (HUVEC), from which we could generate a large amount of fresh primary EC. Thus, we avoided the need to passage, freeze/thaw, or culture the EC for a prolonged period prior to treatment and analysis, factors that could affect behaviour and response to cytokines (Liao et al., 2014; Ohori et al., 2021). However, it should be acknowledged that this EC type is foetal, rather than adult, and various differences have been reported between the two, including the transcriptional response to TNF (Viemann et al., 2006). Comparisons of microarray data for human dermal microvascular EC (HMEC1) and HUVEC have shown that around half of the TNF induced changes were specific for only one or other of these EC types (Viemann *et al*., 2006). However, key genes that were highlighted as only up regulated by TNF in HMEC1, but not HUVEC (e.g., *IL1B*, *DUSP6*, *OAS1*, *CLDN1*, *CD70*, *MMP12*), were classified as up regulated in our HUVEC dataset - potentially indicating that other factors, such as the sensitivity of EC types to passage in culture or freeze thraw cycles could influence response, as opposed to core characteristics of the EC types *per se*. Indeed, other studies show more similar characteristics between EC types, such as the response to shear stress exposure, which was largely comparable between human adult aortic cells and HUVEC (Maurya et al., 2021).

It should also be noted that EC in our study are not cultured under flow, or together with other cell types found in the normal microenvironment, both factors that can affect *in vitro* gene expression (Afshar et al., 2023; Helle et al., 2021; Heydarkhan-Hagvall et al., 2003; Nakajima and Mochizuki, 2017). Exposure to different levels of flow *in vitro* can modify the EC response to TNF (Sheikh et al., 2003) and thus, our data may be more representative of EC responses in low, rather than high, shear exposed vessels. Finally, although under steady-state conditions protein expression is highly dependent on mRNA level (Liu et al., 2016), the relationship between the two after state transitions, such as those induced by TNF, is subject to time dependent processes, such as maturation, export and translation of mRNA (Liu *et al*., 2016). Thus, there will be a delay between transcriptional changes and the associated protein level increase or decrease. Although extensive, our dataset does not provide a comprehensive overview of all aspects of the TNF response; the release of stored and secreted immediate responders, e.g., P-selectin and Von Willebrand factor (Metcalf et al., 2008), and processes such as protein phosphorylation and nuclear translocation are not measured.

## METHODS

### LEAD CONTACT

Further information and requests for resources and reagents should be directed to and will be fulfilled by the Lead Contact: Dr. Lynn Marie Butler.

### MATERIALS AVAILABILITY

This study did not generate new unique reagents.

### DATA AND CODE AVAILABILITY

The data generated by this study is publicly available. Any additional information required to reanalyse the data reported in this paper is available from the lead contact upon request.

### EXPERIMENTAL MODEL AND SUBJECT DETAILS

#### Isolation, culture, and sex-determination of human umbilical vein endothelial cells

Human umbilical vein endothelial cells (EC) were isolated from human umbilical cords, collected from Karolinska University Hospital, Stockholm, Sweden, as described(Cooke et al., 1993). Ethical approval was granted by *Regionala etikprövningsnämnden i Stockholm* (2015/1294-31/2). EC were cultured in Medium M199, supplemented with 10% fetal bovine serum, 10 ml/l penicillin-Streptomycin, 2.5 mg/l Amphotericin B (all ThermoFisher, Gibco), 1 mg/l hydrocortisone and 1 µg/l and human epidermal growth factor (hEGF) (both Merck). To determine EC sex, transcripts encoding *Ubiquitously Transcribed Tetratricopeptide Repeat Containing, Y-Linked* (*UTY*) were measured by qPCR. Cell lysis and cDNA generation was performed using the 2-Step Fast-Cells-to-CT- Kit (Invitrogen, ThermoFisher) according to their protocols. qPCR was performed using TaqMan Fast Universal PCR mix and target (UTY) primer conjugated to FAM-probe (Hs01076483, ThermoFisher) with 18s rRNA primer (4319413E conjugated to VIC probe, ThermoFisher) as endogenous control. qPCR was performed using a RealTime PCR LightCycler 96 ® system (Roche Life Sciences). EC positive for *UTY* expression were classified male, and those negative classified female. 5-6 sex-matched biological replicates were pooled to create each sample set (33 donors in total). Sample sets were A, B, C (annotated male) and D, E, F (annotated female). Following sequencing, a low level of Y-linked transcripts were detected in sample set F, indicating an incorrect annotation of one of the constituent donors as female. Thus, this sample was excluded from any subsequent sex-based analysis, but included in non sex-split analyses.

#### EC treatment, RNA isolation and sequencing

Pooled HUVEC sample sets were grown to confluence before treatment with or without recombinant tumour necrosis factor alpha (TNF; 10 ng/mL) (ThermoFisher) in cell culture medium. EC were lysed at 0.5, 1, 2, 3, 4, 6, 8, 12, 24, 36, 48 or 72 hours post-stimulation, using RLT lysis buffer from the RNAeasy mini kit (Qiagen). RNA isolation and purification was performed using the RNAeasy mini kit (Qiagen). RNA concentration was measured using Nanodrop 2000 spectrophotometer and RNA integrity number (RIN) determined using Agilent 2100 Bioanalyzer (RIN>9 required for inclusion). Library preparation and RNA sequencing was performed by the National Genomics Infrastructure Sweden (NGI) using Illumina stranded TruSeq poly-A selection kit and Illumina NovaSeq6000S (4 lanes, 2x 150bp reads, incl 2Xp kits). The data was processed using demultiplexing. Data storage and initial analyses were performed using server sided computation supplied by the Swedish National Infrastructure for Computing (SNIC). Genome assembly used for sequence alignment: Homo_sapiens.GRCh38.dna.primary_assembly.fa and annotated using Homo_sapiens.GRCh38.96.gtf. Sequence alignment was carried out using STAR/2.5.3a. Gene mapping has been carried out using subread/1.5.2 and the module feature counts. Transcript mapping carried out using Salmon/0.9.1.

### QUANTIFICATION AND STATISTICAL ANALYSIS

#### Gene centric analysis and normalization

Raw gene counts (Table S3) were normalized to TPM (transcripts per kilobase per million reads mapped). TPM values were used to map gene count comparisons between samples across the same groups, with groups being defined by treatment type (“Untreated”, “TNF stimulated”), sex (“female”, “male”) and by time point (hours: 0.5, 1, 2, 4, 6, 8, 12, 24, 36, 48, 72) (“Gene centric differential expression analysis”). To identify genes affected by TNF treatment, gene centric differential expression analysis was performed between TNF treated sample and the matched control.

The analysis was executed by comparing sex-, time- and donor-matched TNF stimulated samples to their respective control and calculating fold change values. Average values were generated by merging same sex replicates for each timepoint or merging all replicate samples for respective timepoints.

#### Data normalization and differential gene expression analysis

We used the “DESeq2” package in R to normalize raw gene expression counts (Love et al., 2014). Normalized expression values were log2 transformed and averaged across biological replicates. To see the clustering of samples, multidimensional scaling (MDS) plots based on overall gene expression for all samples including replicates and averaged expression across biological replicates were created.

To further identify genes affected by TNF treatment, the “DESeq2” R package (Love *et al*., 2014) was used to perform differential expression analysis between treated and untreated samples for each time point, with female and male samples analysed separately. In total, comparisons for 11 time points were performed. Differential expression analysis was also performed for comparisons between sexes at each time point, between treated and untreated samples. We used the DESeq2 time series design to identify genes with different expression at any time point between treated and untreated samples for female and male. In this case, the full model is represented by ∼treatment + time+ treatment:time and the reduced model by ∼treatment+time. In all differential expression comparisons, genes with adjusted P-value <0.05 and an absolute fold change of log2 >1 were classified as differentially expressed genes (DEGs) and total DESeq count below 10 across all samples, were removed from the DEG analysis.

#### Further classification of positive and negative profiles

To identify genes directly regulated by TNF, independent of change in base line, we created subgroups defined by their reaction profile based on the previous analysis. Genes with low expression, i.e., with a maximal TPM values were excluded from DEG analysis. DEGs were clustered into positive or negative regulation profiles. Those that fell into both categories, at different timepoints or in different sexes, were excluded from the analysis. To identify genes that are temporally differentially expressed and remove noise, for inclusion DEGs must have been classified as such at two or more sequential timepoints. We defined profiles for negative DEGs as “down regulated” (where TNF reduced gene expression from a stable baseline), or “temporally delayed”, “inhibited” and “other” (where baseline changes could drive differential expression) (Figure S1 D i-iii). Down regulated profiles expressed a low variation (coefficient of variation (CV) < 0.3) across the untreated samples and a high variation across the TNF-treated samples (CV > 0.3), as well as a log2 fold change of <0.6 in the control and a minimum log2 fold change of > 0.7 in the TNF treated samples. Temporally delayed DEGs were defined as having a high variation within the control (CV > 0.3) and a high variation within the TNF treated samples (CV > 0.3), as well as a minimum log2 fold change of > 0.7 in TNF treated samples. Inhibited DEGs express have a high variation within the control (CV > 0.3) and a low variation within the TNF treated samples (CV < 0.3), as well as a minimum log2 fold change of > 0.7 within the control and a maximum log2 fold change of < 0.6 in the TNF treated samples. DEGs that did not fall into any of the of three described categories were defined as “other”.

Classification of positive DEGs was performed using exclusion criteria, excluding samples with decreasing baseline and stable TNF expression. Excluded samples show a low variation within TNF treated samples (CV <0.3) and a high variation within the baseline samples (CV >0.3) concurring with a transcriptional decrease in fold change (log 2 FC < 0) compared to the initial timepoint (0.5 h) (Figure S1 C i, ii).

#### Gene co-expression network construction

A gene co-expression network was constructed using the WGCNA package in R (Langfelder and Horvath, 2008). After filtering of low expressed genes (total DESeq2 counts over the time course < 10) and genes with low variation across all samples (CV >0.2), 12 428 genes were remained for further analysis.

The appropriate soft-thresholding power was selected by applying “pickSoftThreshold” function with parameter “networkType” set to signed hybrid. Then the correlation network adjacency matrix was calculated using selected soft thresholding of 12 and with parameter “networkType” set to signed hybrid.

The adjacency matrix was turned into topological overlap (TOM) and the corresponding dissimilarity (dissTOM) was calculated. Finally, average linkage hierarchical clustering was applied according to the dissTOM value and gene modules were identified using dynamic tree cut algorithm with the minimum module size of 30 genes. Module eigengenes (MEs) representing the first principal component of each expression module were calculated for each module.

#### Module visualization

The profiles of each module were visualized using a heatmap of scaled gene expression profiles. Additionally, plots displaying the expression of each gene within module and the expression of its eigengene were produced (Langfelder and Horvath, 2008), which returns the gene in each module with the highest connectivity.

#### Functional enrichment of co-expression modules and DEGs

Gene ontology (GO) functional annotation and KEGG / Reactome pathways enrichment analysis of modules was performed using hypergeometric test “hyperGTest” from “GOstats” package in R. For sets of DEGs obtained from different comparisons we performed gene set enrichment analysis (GSEA) on the log fold change from DESeq2 to identify Reactome gene sets enriched for differentially expressed genes.

The Gene Ontology Consortium (Ashburner et al., 2000a) and PANTHER classification resource (Mi, 2019) were used to identify overrepresented terms in gene lists from the GO ontology (release date 2022-07-01) databases. Plots of GO terms were created using the R package clusterProfiler (Wu et al., 2021).

#### Additional statistical analyses and website development

Graphs were created using the packages ggplot2 (Wickham, 2016) and base R(Team, 2022) and GraphPad Prism. Temporal graphs for Figures 3D, 4, S3 were created by using our website tool. Circle plots were created by using the R package circlize (Gu et al., 2014) and PubMed lookups were performed using the R package easyPubMed by Damiano Fantini (Fantini, 2019) (date of lookup 01.03.2023). Figures were assembled using Affinity Designer and Adobe Illustrator.

The following additional R packages were used for mapping, normalization, clustering, and display of the data: readR, dplyr, data. table, matrixStats, NormExpression, edgeR, viridis, RcolorBrewer.

Further statistical analyses and website implementation were performed in RStudio (R version 4.0.3) using shinyapps. The following packages were used: shiny (Chang et al., 2022), shinyjs, gplot, ggplot2, DT, plotly and ggthemes.

The website is available here: https://butlerlab.shinyapps.io/temporal_TNF_response/

#### Website and data availability resource

Average Gene TPM and DEG values (available as log2, log10 or decimal) for each timepoint are available in Table S1, Tab 1 and 2. Raw gene counts are available in Table S3. TPM, DESeq2 and DEG values for each individual donor pool, and data for modules can be downloaded from https://butlerlab.shinyapps.io/temporal_TNF_response/. All gene expression profile plots and data for all genes can be downloaded directly from the website. The data is available as TPM or DEG fold change values and can be split by sex and filtered by observation timeframes of interest.

## AUTHOR CONTRIBUTIONS

Conceptualisation: LMB, ECS, PD. Methodology: LMB, ECS, PD. Formal analysis: ECS, TB, PD, PH, MLK. Investigation: LMB, ECS, PD, TB, PH, MLK. Writing – Original Draft: ECS, LMB, PD. Writing – Review & Editing: All. Website development: ECS, PD. Visualisation: ECS, TB, LMB, PD. Supervision: LMB, PD, MLK. Resources: LMB, JO, MLK. Funding Acquisition: LMB, JO, MLK.

## ACKNOWLEDGEMENTS

Funding was granted to LMB and MLK from NCMM, Oslo University, Norway, to LMB from Hjärt Lungfonden (20170759, 20170537, 20200544) and the Swedish Research Council (2019-01493), and to JO from Stockholm County Council (SLL 2017-0842). Computations and data handling were enabled by resources provided by the Swedish National Infrastructure for Computing (SNIC), partially funded by the Swedish Research Council through grant agreement no. 2018-05973. TB, PHH, and MLK are supported by the Norwegian Research Council, Helse Sør-Øst, and University of Oslo through the Centre for Molecular Medicine Norway (187615).

---

**Figure S1.**
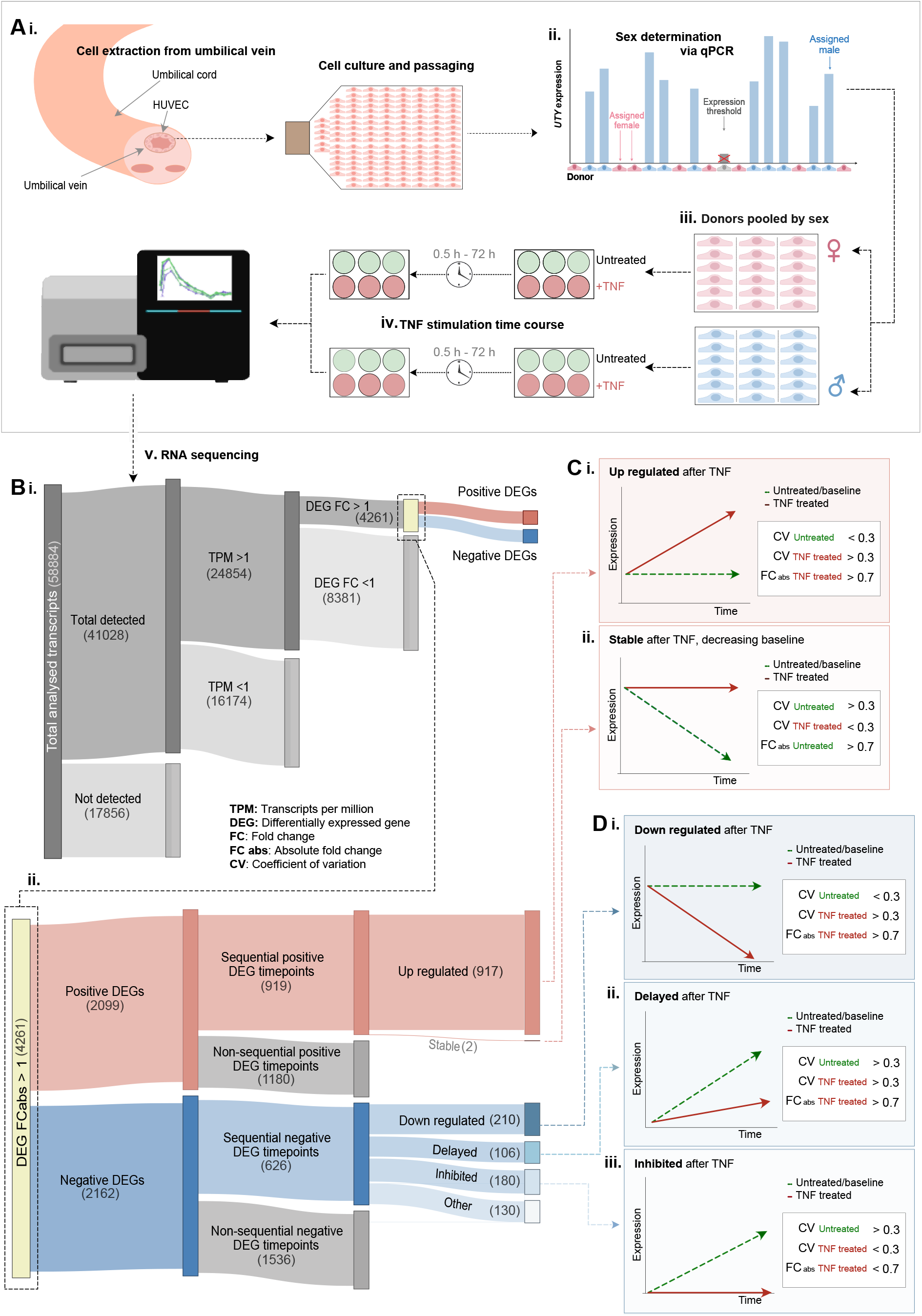
Experimental design and gene classifications. (**A**) Overview of the experimental procedure. (i) Endothelial cells were extracted from human umbilical veins and cultured to confluency before (ii) qPCR for Y-chromosome gene *UTY* was used to identify male vs. female donors. (iii) Cells were pooled into sex-matched sample sets, grown to confluency and (iv) stimulated with tumour necrosis factor alpha (TNF; 10 ng/mL), before RNA extraction at 0.5, 1, 2, 4, 6, 8, 12, 24, 36, 48 or 72 hours post stimulation, followed by (v) RNA sequencing analysis. (**B**) Sankey plots displaying (i) total number of genes detected and classified as differentially expressed genes (DEG) (ii) numbers of positive and negative DEG, and the subsequent classification as Positive DEGs as: (i) *up regulated* by TNF from a stable baseline expression in control EC, or (ii) *stable* on the background of reduced baseline expression in control EC over time. (**D**) Negative DEGs as: (i) *down regulated* by TNF from a stable baseline expression in control EC, or (ii) *delayed* or (iii) *inhibited* by TNF, where baseline gene expression in control EC increases over time, but this change either occurs later or is not observed in TNF treated EC, respectively. Genes that did not fall into these groups were categorised as ‘other’.

**Figure S2.**
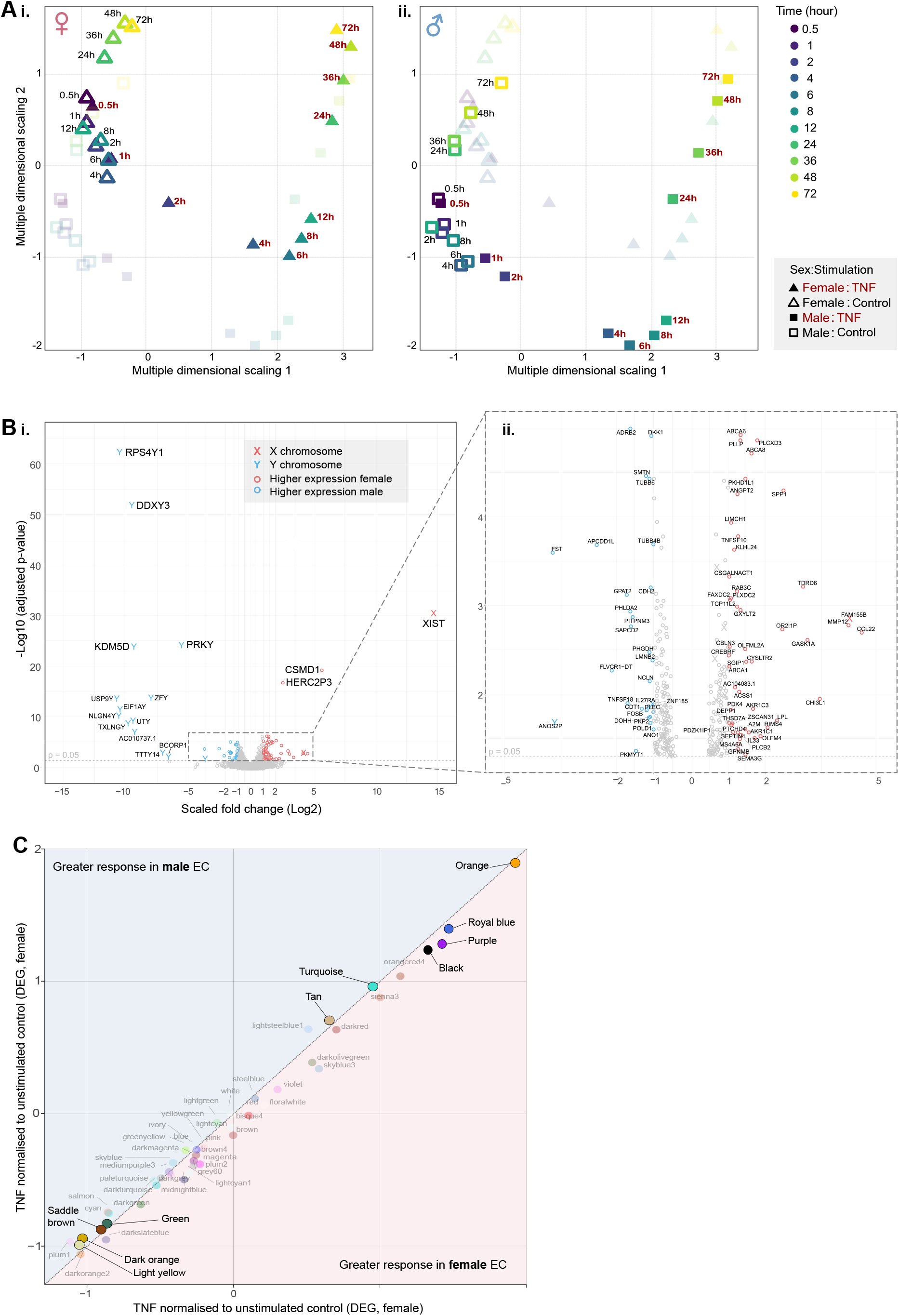
Sex-based comparison of global gene expression profiles. Human umbilical vein endothelial cells (EC, male n=3, female n=2) were treated with or without tumour necrosis factor alpha (TNF) and harvested at 0.5, 1, 2, 4, 6, 8, 12, 24, 36, 48 or 72 hrs, before RNAseq analysis. (**A**) Multidimensional scaling plot for control or TNF treated (i) female or (ii) male EC, at all analysed time points. (**B**) Volcano plot displaying differentially expressed genes between male and female EC under baseline (unstimulated control) conditions, with X- and Y-chromosomal genes highlighted (**C**) Scatter plot comparing normalised average expression values between male and female samples for each WGCNA module.

**Figure S3.**
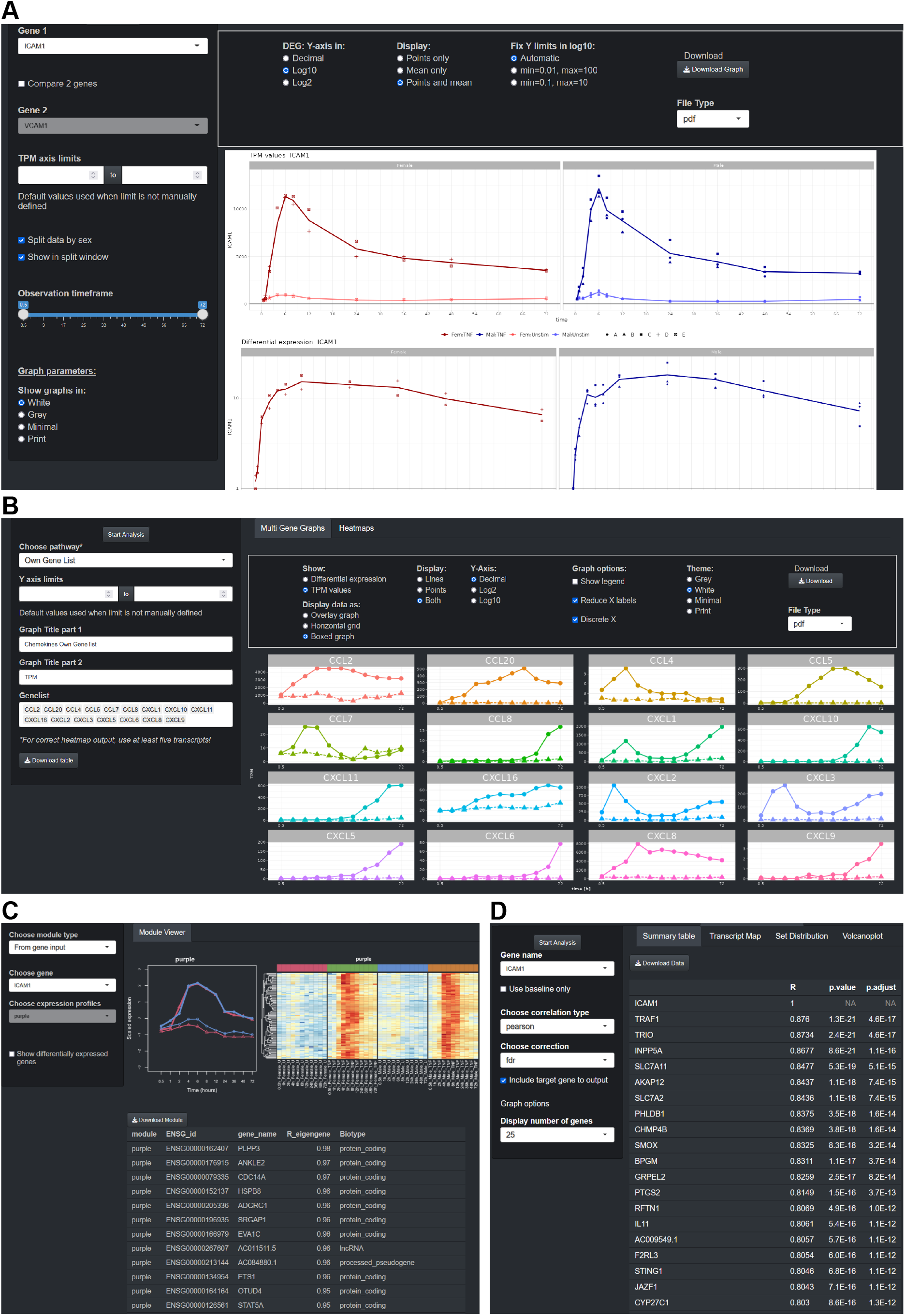
Website resource examples. All data generated in this study is available on https://butlerlab.shinyapps.io/temporal_TNF_response/. Selected features include: (**A**) data viewer for temporal gene expression over time (displayed as absolute values and relative differential expression), (**B**) data plot generator to be used either with user defined gene lists, or predefined gene categories, selected from the dropdown menu (e.g., ‘leukocyte recruitment’), (**C**) weighted network correlation analysis data section and module look up tool for any given input gene, (**D**) Expression similarity tool that can identify genes with the highest correlation to any given input gene across the dataset. Data plots can be downloaded as vector-images.

